# Glycine fermentation by *C. difficile* promotes virulence, spore formation, and is induced by host cathelicidin

**DOI:** 10.1101/2023.06.30.547236

**Authors:** Arshad Rizvi, Germán Vargas-Cuebas, Adrianne N. Edwards, Michael A. DiCandia, Zavier A. Carter, Cheyenne D. Lee, Marcos P. Monteiro, Shonna M. McBride

## Abstract

The amino acid glycine is enriched in the dysbiotic gut and is suspected to contribute to *Clostridioides difficile* infection. We hypothesized that the use of glycine as an energy source contributes to colonization of the intestine and pathogenesis of *C. difficile*. To test this hypothesis, we deleted the glycine reductase genes *grdAB*, rendering *C. difficile* unable to ferment glycine, and investigated the impact on growth and pathogenesis. We found that the *grd* pathway promoted growth, toxin production, sporulation, and pathogenesis of *C. difficile* in the hamster model of disease. Further, we determined that the *grd* locus is regulated by host cathelicidin (LL-37) and the cathelicidin-responsive regulator, ClnR, indicating that the host peptide signals to control glycine catabolism. The induction of glycine fermentation by LL-37 demonstrates a direct link between the host immune response and the bacterial reactions of toxin production and spore formation.

## INTRODUCTION

*C. difficile* is an anaerobic, Gram-positive, spore-forming gastrointestinal pathogen that causes severe diarrheal disease which can lead to death (Sandhu and McBride, 2018). To cause disease, *C. difficile* spores must be ingested and germinate in the intestine upon exposure to bile salts and glycine (Sorg and Sonenshein 2008). Upon the outgrowth of spores into vegetative cells, *C. difficile* colonizes the colon, using the available nutrients to replicate within the host. As the population increases and nutrients become scarce, *C. difficile* produces toxins (TcdA and TcdB), releasing additional nutrients and further altering the metabolic environment (Fletcher *et al*., 2021). In turn, host inflammation is induced, resulting in the release of immune effectors and the characteristic pathologies associated with *C. difficile* infections (CDI) (Abt *et al*., 2016).

*C. difficile* infections are typically preceded by intestinal dysbiosis, resulting in enrichment of amino acids that *C. difficile* preferentially uses to generate energy (Battaglioli *et al*., 2018). *C. difficile* catabolizes amino acids through fermentation pathways known as Stickland reactions, which involve the coupled oxidation and reduction of amino acid pairs (Stickland, 1934; Stickland, 1935; Hofmann *et al*., 2018). The amino acid glycine is a known Stickland substrate in *C. difficile*, in addition to its role as a spore co-germinant (Jackson *et al*., 2006; Sorg and Sonenshein, 2008; Bouillaut *et al*., 2013). Glycine is catabolized by the glycine reductase (GR) pathway, which is encoded by the *grd* genes (Andreesen, 2004). While glycine fermentation is known to occur in *C. difficile*, the importance of this pathway to *C. difficile* development and pathogenesis is not known (Bouillaut *et al*., 2013; Hofmann *et al*., 2018; Johnstone and Self, 2022).

Despite the presence of glycine reductase pathways in several anaerobic pathogens, the association of glycine catabolism with virulence and cell physiology has not been explored. In previous work, we identified the LL-37-responsive regulator, ClnR, as a putative regulator of glycine catabolism (Woods *et al*., 2018). In this study, we investigated the role of glycine Stickland metabolism in the growth and virulence of *C. difficile* and the regulation of this metabolic pathway by the host inflammatory peptide, LL-37. We present *in vitro* and *in vivo* evidence that indicate glycine is an important metabolic pathway for *C. difficile* pathogenesis and spore formation. Additionally, we provide evidence that the induction of glycine catabolism occurs in response to host inflammation through the regulator ClnR.

## RESULTS

### The host peptide LL-37 and glycine promote glycine catabolism and enhance *C. difficile* growth

We previously showed that *C. difficile* regulates the expression of metabolic pathways in response to the host peptide LL-37, including the glycine fermentation pathway (Woods, Edwards et al. 2018). From the increased expression of the *grd* operon previously observed with LL-37, we hypothesized that in the presence of LL-37, *C. difficile* would use glycine more efficiently as an energy source. To test this hypothesis, we generated a deletion mutant that lacks the catalytic subunits of the glycine reductase complex, GrdA and GrdB (**Fig S1**; Δ*grdAB*, MC1576), rendering *C. difficile* unable to use glycine for energy through Stickland fermentation. The deletion of *grdAB* was verified as the only genetic modification in this strain by whole genome sequencing (NCBI SRA#: SAMN23038927).

To determine the impact of the *grdAB* mutant on glycine catabolism, we assessed growth in minimal media (CDMM) with or without glycine, LL-37, or both. No difference in growth was observed between the parent strain 630Δ*erm* (WT) or the *grdAB* mutant during exponential phase in any condition (**Fig 1A**). However, upon transition to stationary phase, the WT achieved a higher cell density than the *grdAB* mutant in all conditions, indicating that glycine catabolism increases in stationary phase. WT growth also increased modestly when supplemented with LL-37, as previously observed (Woods *et al*., 2018). However, when LL-37 and glycine were both added, growth of the WT strain was further enhanced, while the *grdAB* mutant growth remained unchanged, indicating that glycine catabolism was enhanced by the presence of LL-37 (**Fig 1A,B)**.

**Figure 1.**
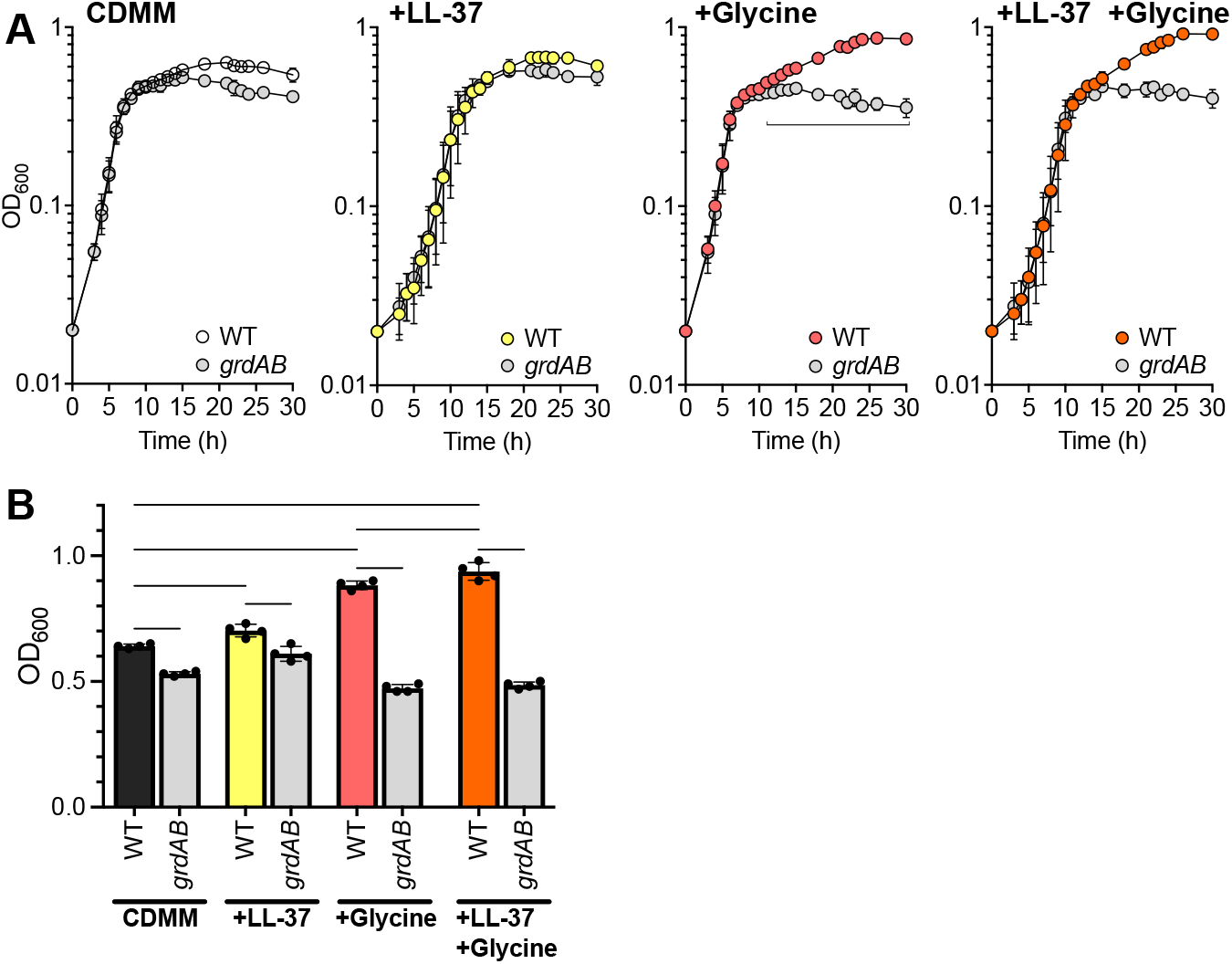
The host peptide LL-37 and glycine catabolism enhance growth of *C. difficile*. A) Active cultures of strain 630Δ*erm* (WT) and the *grdAB* mutant (MC1576) were grown in CDMM, or CDMM supplemented with LL-37 (0.5 µg/ml), glycine (30 mM), or both LL-37 and glycine. Graphs are plotted as the means +/- SD from four independent replicates. Data were analyzed by Student’s *t* test comparing the mean values of *grdAB* to WT at each time point. B) Maximum cell density of WT or *grdAB* mutant cultures in different conditions. The mean value and SD of four replicates for each condition and strain are shown. Data were analyzed by 2-way ANOVA followed by Tukey’s multiple comparisons test. ^*^, P < 0.05; ^**^ P < 0.01; ^***^, P < 0.001; ^****^ P < 0.0001.

### Glycine Stickland metabolism is required for efficient sporulation and delays spore germination

To determine if glycine catabolism affects spore formation, we examined sporulation of the WT and *grdAB* strains after growth on sporulation agar. The production of ethanol-resistant spores was reduced five-fold in the *grdAB* mutant (WT, 16.5 ± 0.8%; *grdAB*, 3.7 ± 0.7%), indicating that glycine catabolism supports efficient sporulation (**Fig 2A,B**). Phase-contrast microscopy of the sporulating cells revealed fewer phase-bright spores formed by the *grdAB* mutant relative to the parent strain, indicating that the *grdAB* mutant was less effective at progressing to fully formed spores.

**Figure 2.**
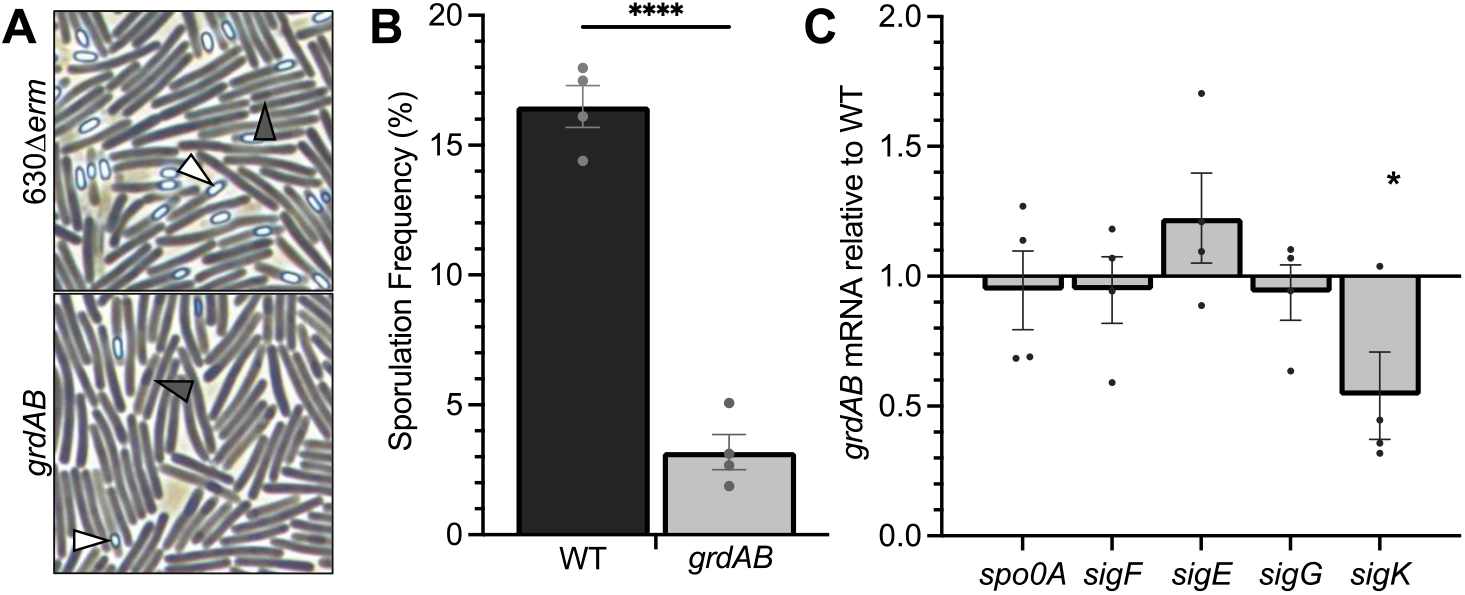
Glycine utilization promotes sporulation and spore development. A) Representative phase-contrast micrographs of 630Δ*erm* and the *grdAB* mutant (MC1576) grown on sporulation agar for 24 h. White arrowheads indicate phase bright spores, and dark arrowheads indicate phase dark spores. B) Ethanol-resistant spore formation for 630Δ*erm* and the *grdAB* mutant. ^****^ P<0.0001 by a two-tailed Student’s *t* test. C) qRT-PCR analysis of transcripts of the major sporulation regulators *spo0A, sigF, sigE, sigG*, and *sigK*. The means and individual values for four biological replicates are shown. ^*^, P<0.05 using the two-tailed Student’s *t* test comparing each transcript in the *grdAB* mutant to the isogenic parent strain.

Spore formation in *C. difficile* is defined by morphological stages that progress with the sequential activation of sporulation sigma factors (Paredes-Sabja *et al*., 2014). To understand how the sporulation program was affected by the inability to catabolize glycine, we compared expression of the sporulation sigma factors in the *grdAB* and parent strain (**Fig 2C**). No major differences were observed between the *grdAB* and WT strains for the expression of early and mid-stage sigma factors, *sigF, sigE*, and *sigG*, but the transcription of *sigK* was significantly decreased in the *grdAB* mutant (**Fig 2C**). SigK activates late-stage mother cell transcription and promotes the progression to phase-bright spores. These data suggest that glycine catabolism is needed for efficient spore formation, particularly activation of *sigK* transcription and the completion of late-stage sporulation processes.

As glycine is a co-germinant for *C. difficile* spores and glycine fermentation is important for spore formation, we investigated the impact of the *grd* pathway on germination. We observed significantly more rapid germination of the *grdAB* mutant spores than the parent strain when spores were exposed to taurocholate in rich medium containing glycine (**Fig 3**). These results suggest that glycine fermentation is important for growth and completion of spore formation, leading to spores that are more sensitive to germinant.

**Figure 3.**
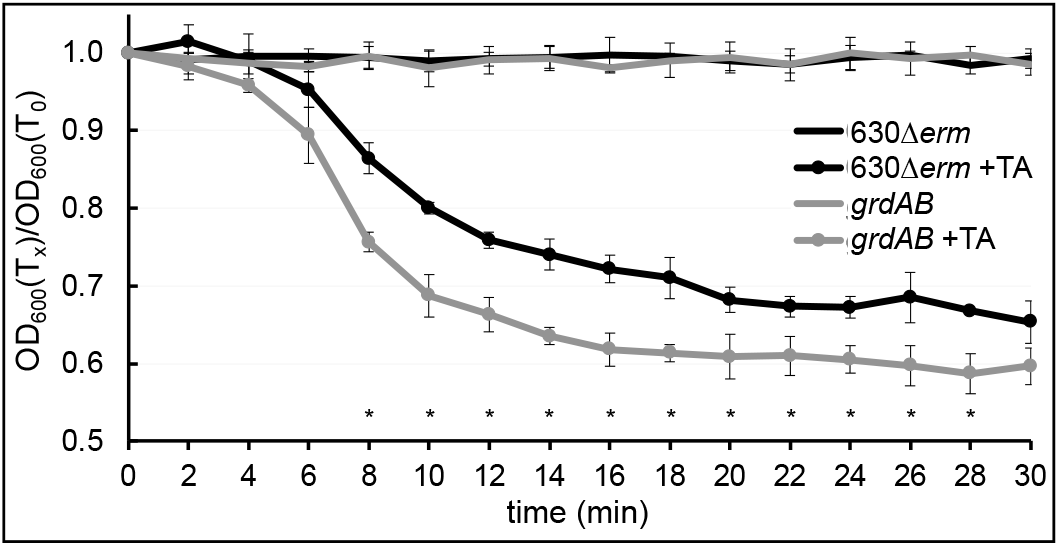
Glycine catabolism impedes spore germination. Purified spores of *C. difficile* 630Δ*erm* (WT) and the *grdAB* mutant (MC1576) were assessed for germination in BHIS +/- 5 mM taurocholate. The optical densities of spore samples were measured and the ratio of the OD_600_ at each timepoint were plotted against the density observed prior to the addition of germinant. The means for four biological replicates are shown. ^*^, P<0.05 using the two-tailed Student’s *t* test comparing each transcript in the *grdAB* mutant to the isogenic parent strain.

### Glycine utilization promotes *C. difficile* pathogenesis

Since growth decreased in the absence of glycine fermentation, we hypothesized that the *grdAB* mutant may be defective for colonization or virulence during growth in the host intestine. To test this, hamsters were inoculated with spores of the parent strain 630Δ*erm* (WT) or *grdAB* mutant and monitored for disease progression. The mean times to morbidity were 48.7 ± 10.7 h for hamsters infected with the WT strain and 70.9 ± 44.2 h for the *grdAB* mutant, with significant differences in median morbidities, indicative of decreased virulence with the mutant strain **(Fig 4A**). High variability in the time to morbidity was observed for *grdAB* mutant infections, with some animals experiencing prolonged infection. To determine if the differences observed in survival times for the WT and *grdAB* mutant were due to reduced colonization, we analyzed the bacterial output in cecal contents of the infected animals. No difference was observed in *C. difficile* carriage between animals infected with the WT or the *grdAB* mutant (**Fig 4B**). The amount of toxins present in the cecal contents at the time of death were also similar in the *grdAB* mutant infected animals, as well as *in vitro* (**Fig 4C, Fig S2**). These data suggest that glycine catabolism does not affect *C. difficile* host colonization or carriage during infection, but does promote virulence and toxin production in the host.

**Figure 4.**
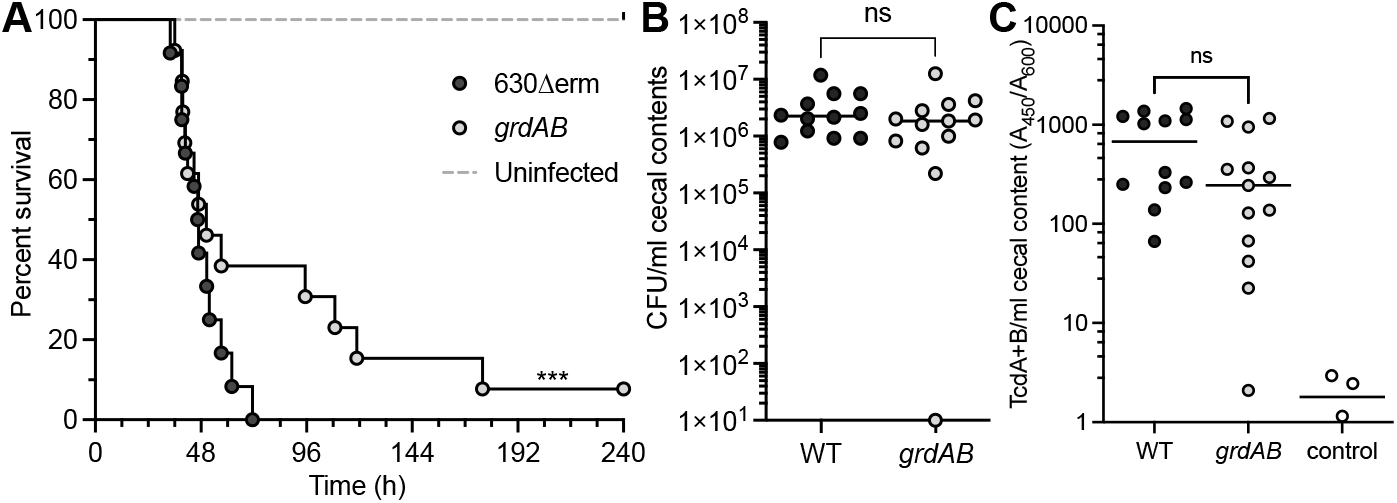
Glycine catabolism promotes virulence. A) Kaplan-Meier survival curve depicting the time to morbidity of Syrian golden hamsters infected with 630Δ*erm* (n = 12) or the g*rdAB* mutant (MC1576; n = 13). The mean times to morbidity were: 630Δ*erm*, 48.7 ± 10.7 h; *grdAB*, 70.9 ± 44.2 h. ^***^, P<0.01, by log-rank test. B) The total *C. difficile* CFU and C) toxin recovered from cecal content post-mortem. Solid lines represent the median value for each strain. ns=not significant by Mann-Whitney test (B) or unpaired *t* test (C).

### LL-37 and glycine increase transcription of the *grd* operon

To better understand the phenotypes associated with glycine fermentation, we investigated transcription and regulation of the *grd* operon. Prior studies identified a promoter upstream of *grdE* (Bouillaut, Self et al. 2013); however, an additional promoter is predicted upstream of *grdX*. We first assessed expression of the *grd* genes by RT-PCR and determined that the *grdX* through *grdD* genes are co-transcribed, indicative of a functional promoter within the region upstream of *grdX* (**Fig S3**).

We next delineated promoter elements within the *grd* region using transcriptional fusions of fragments to an alkaline phosphatase (AP) reporter **(Fig 5A**). *C. difficile* containing promoter::*phoZ* fusions were grown in CDMM with and without LL-37 or glycine, and culture samples were assessed for AP activity. Only the fragment upstream of *grdX* (F1::*phoZ*) generated AP activity under the conditions tested. Further, the F1/P*grdX::phoZ* fusion demonstrated greater reporter activity when cells were grown with LL-37, glycine, and LL-37+glycine, than with CDMM alone, indicating that the P*grdX* promoter is responsible for activation of *grd* expression under these conditions.

**Figure 5.**
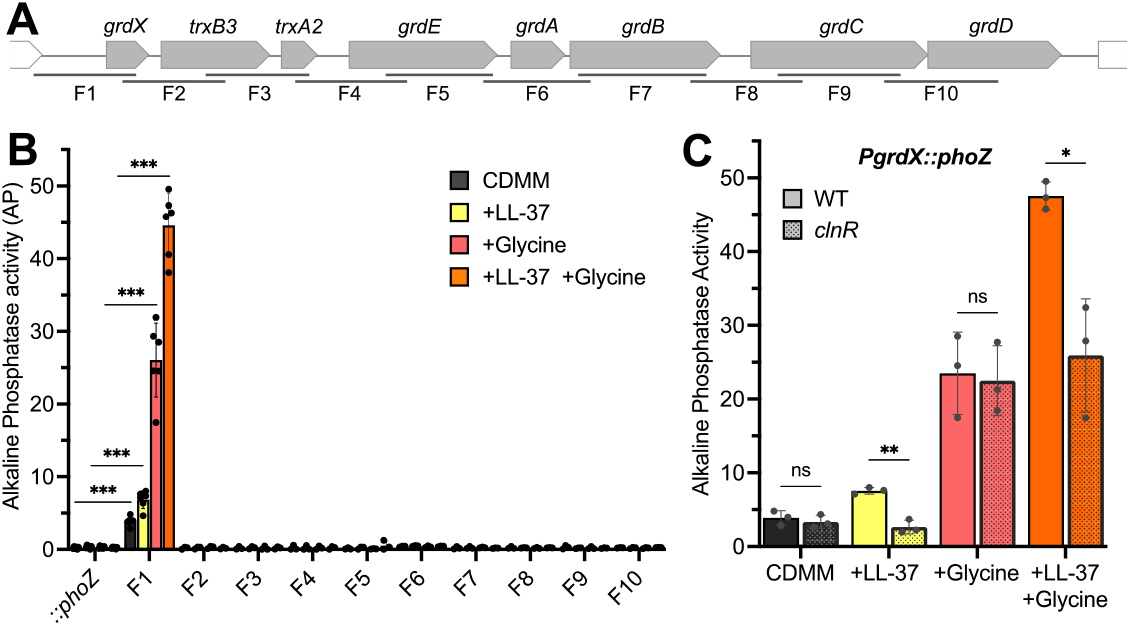
LL-37 activates transcription of glycine catabolism from the *grdX* promoter. A) Schematic of DNA fragments (F1-F10) of the *grd* region transcriptionally fused to *phoZ*. B) Alkaline phosphatase (AP) activity detected from *C. difficile* strain 630Δ*erm* carrying different *grd* region fragments fused to *phoZ* and grown in CDMM + 2 µg ml^-1^ thiamphenicol with or without LL-37 and glycine, as indicated. Statistical significance was assessed by 1-way ANOVA within each growth condition compared to the promoterless *::phoZ* control. C) AP activity from the F1 promoter (P*grdX::phoZ*) in WT (MC1649) or the *clnR* mutant (MC1650). Statistical significance was assessed by Student’s *t* test, comparing activity in the WT and *clnR* mutant from each condition. The mean values with standard deviation of a minimum of three biological replicates are shown.

In *C. difficile*, the transcriptional regulator ClnR binds to LL-37 to regulate transcription of hundreds of genes involved in metabolism, virulence, and growth (Woods, Edwards et al. 2018). To determine if ClnR is responsible for regulating P*grdX* transcription in LL-37, we analyzed the activity of the P*grdX* (F1) fusion in a *clnR* mutant, compared to the WT strain (**Fig 5B**). The *clnR* mutant yielded less P*grdX* reporter activity than the parent strain when LL-37 or LL-37 + glycine were present, implicating ClnR in the synergistic activation of glycine catabolism by LL-37. Of note, there is a predicted glycine-responsive riboswitch located immediately upstream of *grdX* (Soutourina *et al*., 2013). P*grdX* reporter activity is increased in the presence of glycine in a ClnR-independent manner (**Fig 5B**), providing the first biological evidence that glycine may directly interact with a riboswitch to activate *grd* operon expression in *C. difficile*.

### ClnR-dependent regulation of P*grdX*

The promoter fusion data indicated that the region upstream of *grdX* contains the promoter involved in activation of expression by glycine and LL-37. To characterize P*grdX* regulation further, we next mapped the transcriptional start sites in P*grdX* by 5’ RACE. Several potential transcriptional start sites (TSS) were identified, with the most represented transcript beginning at 297 nt upstream of *grdX*. This TSS was previously identified by 5’-end RNA-seq (Soutourina *et al*., 2013) and is downstream of a predicted SigA-dependent promoter (**Fig S4A**). Additional TSS were identified, but no promoter elements were apparent nearby. These additional transcriptional start sites were mapped within areas upstream of *grdX* that have a high degree of structural complexity, including the predicted glycine riboswitch and an inverted repeat.

We previously demonstrated that ClnR binds to DNA to regulate expression of multiple target genes, but ClnR binding to the P*grdX* region was not investigated (Woods, Edwards et al. 2018). Based on the *clnR* and LL-37-dependent expression data, we hypothesized that ClnR directly regulates the *grd* operon by binding upstream of *grdX*. To test this hypothesis, we performed electrophoretic mobility shift assays (EMSA) to assess ClnR binding within this region. We designed DNA probes encompassing either the predicted SigA promoter region upstream of the mapped transcriptional start site at -297 nt (P*grdX-1*) or the region immediately upstream of the *grdX* coding region which contains a near-perfect inverted repeat and another mapped transcriptional start site at -23 nt (P*grdX-2*) (**Fig S4A**). We observed ClnR binding to P*grdX-1* beginning at 4 μM ClnR, resulting in an apparent *K*_D_ of 5.3 ± 0.4 µM (**Fig S4B)**. While ClnR exhibited some affinity to P*grdX-2*, a full shift was not observed at high ClnR concentrations (**Fig S4C**). Competition assays with unlabeled specific P*grdX* and nonspecific P*spo0A* revealed that ClnR-binding to both DNA probes was not highly specific (**Fig S4B,C**). These results indicate that ClnR can bind directly to the P*grdX-1* promoter; however, the binding affinity and specificity of these interactions *in vitro* were poor under the conditions tested.

## DISCUSSION

In this study, we investigated the role of glycine Stickland metabolism in *C. difficile* growth and pathogenesis. We demonstrated that the use of glycine as an energy source is important for *C. difficile* growth, and glycine utilization is further improved by the host peptide LL-37 and glycine availability (**Fig 1**). In addition, we found that the inability to utilize glycine for energy decreased pathogenesis and sporulation of *C. difficile*, and resulted in faster spore germination (**Fig 2,3,4**).

Stickland metabolism is an important method of energy production for *C. difficile*, though the investigation of individual Stickland metabolites on growth and physiology thus far has focused primarily on proline (Bouillaut *et al*., 2013; Neumann-Schaal *et al*., 2019; Bouillaut *et al*., 2019; Johnstone and Self, 2022). Results from other groups have demonstrated that glycine and the *grd* genes of *C. difficile* are important for enhancing growth, despite differences in the strains investigated (Karasawa *et al*., 1995; Bouillaut *et al*., 2013; Collery *et al*., 2017; Johnstone and Self, 2022). However, glycine catabolism is secondary to preferred amino acids such as proline, which represses *grd* expression (Bouillaut *et al*., 2013).

Although glycine is not utilized first among the Stickland substrates by *C. difficile* (Neumann-Schaal *et al*., 2015), our results show that the ability to catabolize glycine is important for expression of the late-stage sporulation regulator, *sigK*, and is vital for proper spore development (**Fig 2)**. But, how glycine signals to regulate sporulation is not apparent. Nutritional regulators that impact sporulation, such as CodY and CcpA, regulate sporulation initiation and are not activated by glycine availability (Antunes *et al*., 2012; Nawrocki *et al*., 2016; Lee *et al*., 2022). The deficiencies observed in spore maturation suggest that glycine may serve as a specific energy source for *C. difficile* spore development, though further investigation is needed to determine how glycine reduction supports spore maturity.

Several regulatory factors are known to impact transcription of the glycine Stickland pathway, including SigH, SigL, CodY, CcpA, Fur, and Rex (Dineen *et al*., 2010; Saujet *et al*., 2011; Antunes *et al*., 2012; Berges *et al*., 2018; Neumann-Schaal *et al*., 2019; Bouillaut *et al*., 2019; Soutourina *et al*., 2020). The induction of glycine catabolism by LL-37 and ClnR are further evidence linking host environmental cues and regulation of *C. difficile* metabolism (Woods *et al*., 2018). The observed binding of the ClnR regulator to the P*grdX* region was not stringent under the *in vitro* conditions tested; however, our attempts to demonstrate LL-37-dependent ClnR binding *in vitro* have been unsuccessful for all targets examined thus far, including P*clnR*. Since ClnR and LL-37 have demonstrated nanomolar affinity by SPR (Woods *et al*., 2018), we surmise that our EMSA conditions are missing an innate requirement for binding. Thus, we cannot currently resolve whether ClnR binds specifically to P*grdX* under other conditions or regulates *grd* expression indirectly.

The production of toxin in *C. difficile* is linked to nutrient availability (Bouillaut *et al*., 2015). The abundance of energy sources such as glucose, cysteine, ethanolamine, branched-chain amino acids, and proline, are known to decrease toxin production (Dineen *et al*., 2007; Karlsson *et al*., 2008; Bouillaut *et al*., 2013; Nawrocki *et al*., 2018). However, animals infected with the *grdAB* mutant yielded similar levels of toxin, indicating that glycine catabolism did not significantly impact toxin production *in vivo* (**Fig 4**). Mutant-infected animals also survived longer than wild-type infected animals, yet, the *grdAB* mutant did not have an observable colonization defect, as evidenced from similar CFU recovered from infected animals. The ability of the *grdAB* mutant to colonize the intestine, but with reduced pathogenesis, is consistent with the observed stationary-phase increase in glycine utilization (**Fig 1**) and suggests that the use of glycine as an energy source is most significant after *C. difficile* achieves high cell densities in the intestine.

Glycine catabolism promotes toxin production and sporulation, and is induced by host LL-37, providing a link between the host immune response, pathogenesis and dormancy. Understanding the mechanisms of *C. difficile* metabolic adaptation to the host and how these pathways impact pathogenesis has the potential to reveal nutritional strategies to combat infection and transmission. Further, the glycine reductase pathway is encoded within several enteric spore formers, including *Clostridium botulinum, Clostridium scindens, Clostridium sporogenes, Clostridium histolytica*, and *Paeniclostridium sordellii*, and potentially affects sporulation and toxin production in these infections.

## ACKNOWLEDGEMENTS

The authors would like to thank the members of the McBride lab for useful feedback and suggestions during the course of this work. This work was supported by the U.S. National Institutes of Health through research grants AI116933 and AI156052 to S.M.M., T32 AI106699 to C.D.L. and G.V.C., F31 DK126467 to G.V.C., and T32 GM008490 to M.A.D. The content of this manuscript is solely the responsibility of the authors and does not necessarily reflect the official views of the National Institutes of Health.

## AUTHOR CONTRIBUTIONS

S.M.M., A.R., A.N.E., G.V.C., M.A.D., C.D.L., and M.P.M. designed and conducted experiments. S.M.M., A.R., A.N.E., C.D.L., G.V.C., and M.A.D. wrote and edited the manuscript.

## MATERIALS AND METHODS

### Strain and plasmid construction

*C. difficile* strain 630 DNA sequence (RT012; Genbank no. NC_009089.1) was used as a template for designing primers and mapping DNA probes and fusions. Cloned PCR fragments were confirmed by Sanger sequencing (Genewiz). The plasmids and strains used in this study are listed in **Table 1**. Details of strain and plasmid construction are shown in the Supplemental Material (**Fig S5**). Oligonucleotides used for PCR and qRT-PCR analyses are listed in **Table 2**.

**Table 1.**
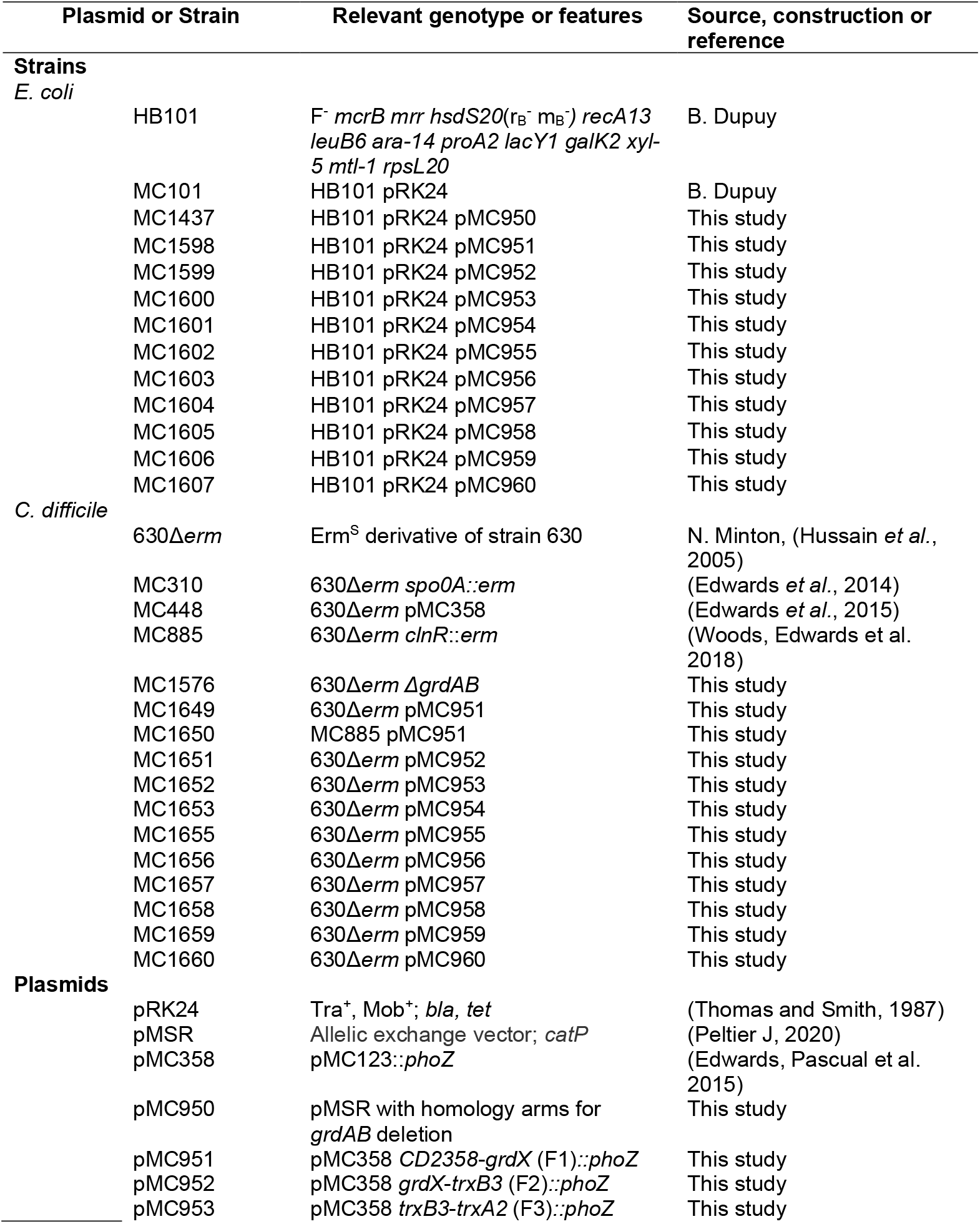
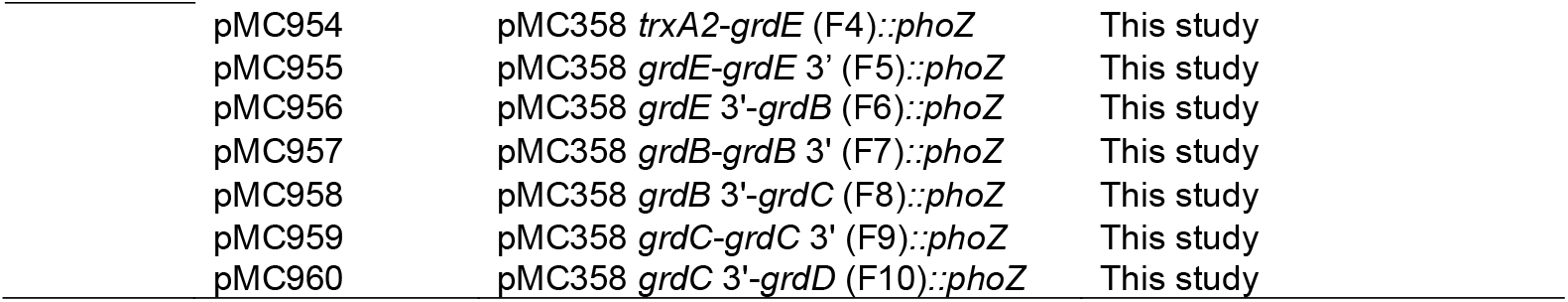
Bacterial Strains and plasmids.

| Plasmid or Strain | Relevant genotype or features | Source, construction or reference |
| --- | --- | --- |
| **Strains** |  |  |
| *E. coli* |  |  |
| HB101 | F<sup>-</sup> *mcrB mrr hsdS20*(r<sub>B</sub><sup>-</sup> m<sub>B</sub><sup>-</sup>) *recA13 leuB6 ara-14 proA2 lacY1 galK2 xyl-5 mtl-1 rpsL20* | B. Dupuy |
| MC101 | HB101 pRK24 | B. Dupuy |
| MC1437 | HB101 pRK24 pMC950 | This study |
| MC1598 | HB101 pRK24 pMC951 | This study |
| MC1599 | HB101 pRK24 pMC952 | This study |
| MC1600 | HB101 pRK24 pMC953 | This study |
| MC1601 | HB101 pRK24 pMC954 | This study |
| MC1602 | HB101 pRK24 pMC955 | This study |
| MC1603 | HB101 pRK24 pMC956 | This study |
| MC1604 | HB101 pRK24 pMC957 | This study |
| MC1605 | HB101 pRK24 pMC958 | This study |
| MC1606 | HB101 pRK24 pMC959 | This study |
| MC1607 | HB101 pRK24 pMC960 | This study |
| *C. difficile* |  |  |
| 630Δ*erm* | Erm<sup>S</sup> derivative of strain 630 | N. Minton, (Hussain *et al.*, 2005) |
| MC310 | 630Δ*erm spo0A::erm* | (Edwards *et al.*, 2014) |
| MC448 | 630Δ*erm* pMC358 | (Edwards *et al.*, 2015) |
| MC885 | 630Δ*erm clnR::erm* | (Woods, Edwards *et al.* 2018) |
| MC1576 | 630Δ*erm ΔgrdAB* | This study |
| MC1649 | 630Δ*erm* pMC951 | This study |
| MC1650 | MC885 pMC951 | This study |
| MC1651 | 630Δ*erm* pMC952 | This study |
| MC1652 | 630Δ*erm* pMC953 | This study |
| MC1653 | 630Δ*erm* pMC954 | This study |
| MC1655 | 630Δ*erm* pMC955 | This study |
| MC1656 | 630Δ*erm* pMC956 | This study |
| MC1657 | 630Δ*erm* pMC957 | This study |
| MC1658 | 630Δ*erm* pMC958 | This study |
| MC1659 | 630Δ*erm* pMC959 | This study |
| MC1660 | 630Δ*erm* pMC960 | This study |
| **Plasmids** |  |  |
| pRK24 | Tra<sup>+</sup>, Mob<sup>+</sup>; *bla*, *tet* | (Thomas and Smith, 1987) |
| pMSR | Allelic exchange vector; *catP* | (Peltier J, 2020) |
| pMC358 | pMC123::*phoZ* | (Edwards, Pascual *et al.* 2015) |
| pMC950 | pMSR with homology arms for *grdAB* deletion | This study |
| pMC951 | pMC358 *CD2358-grdX* (F1)::*phoZ* | This study |
| pMC952 | pMC358 *grdX-trxB3* (F2)::*phoZ* | This study |
| pMC953 | pMC358 *trxB3-trxA2* (F3)::*phoZ* | This study |
| pMC954 | pMC358 *trxA2-grdE* (F4)::*phoZ* | This study |
| pMC955 | pMC358 *grdE-grdE* 3' (F5)::*phoZ* | This study |
| pMC956 | pMC358 *grdE* 3'-*grdB* (F6)::*phoZ* | This study |
| pMC957 | pMC358 *grdB-grdB* 3' (F7)::*phoZ* | This study |
| pMC958 | pMC358 *grdB* 3'-*grdC* (F8)::*phoZ* | This study |
| pMC959 | pMC358 *grdC-grdC* 3' (F9)::*phoZ* | This study |
| pMC960 | pMC358 *grdC* 3'-*grdD* (F10)::*phoZ* | This study |

**Table 2.**
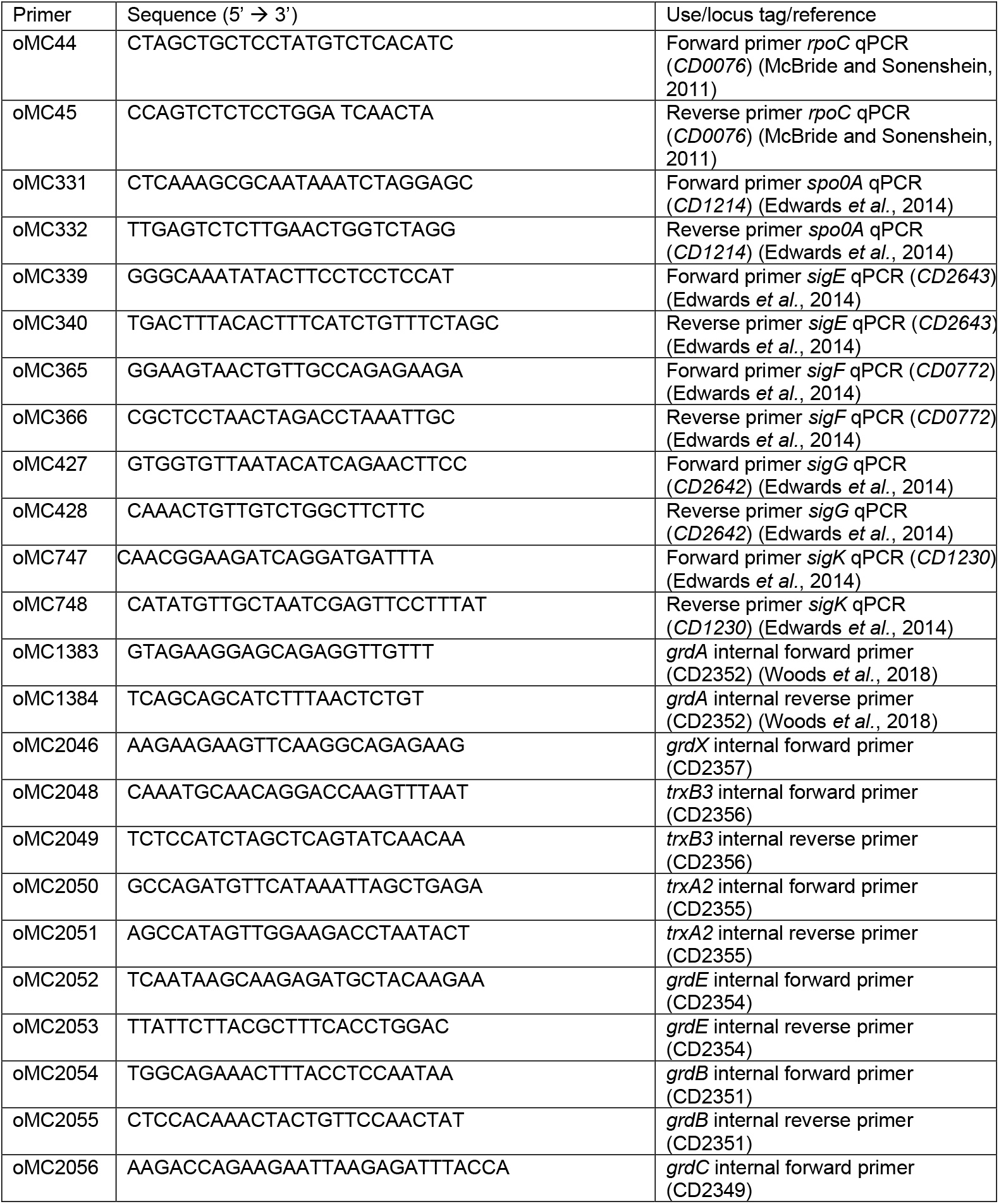
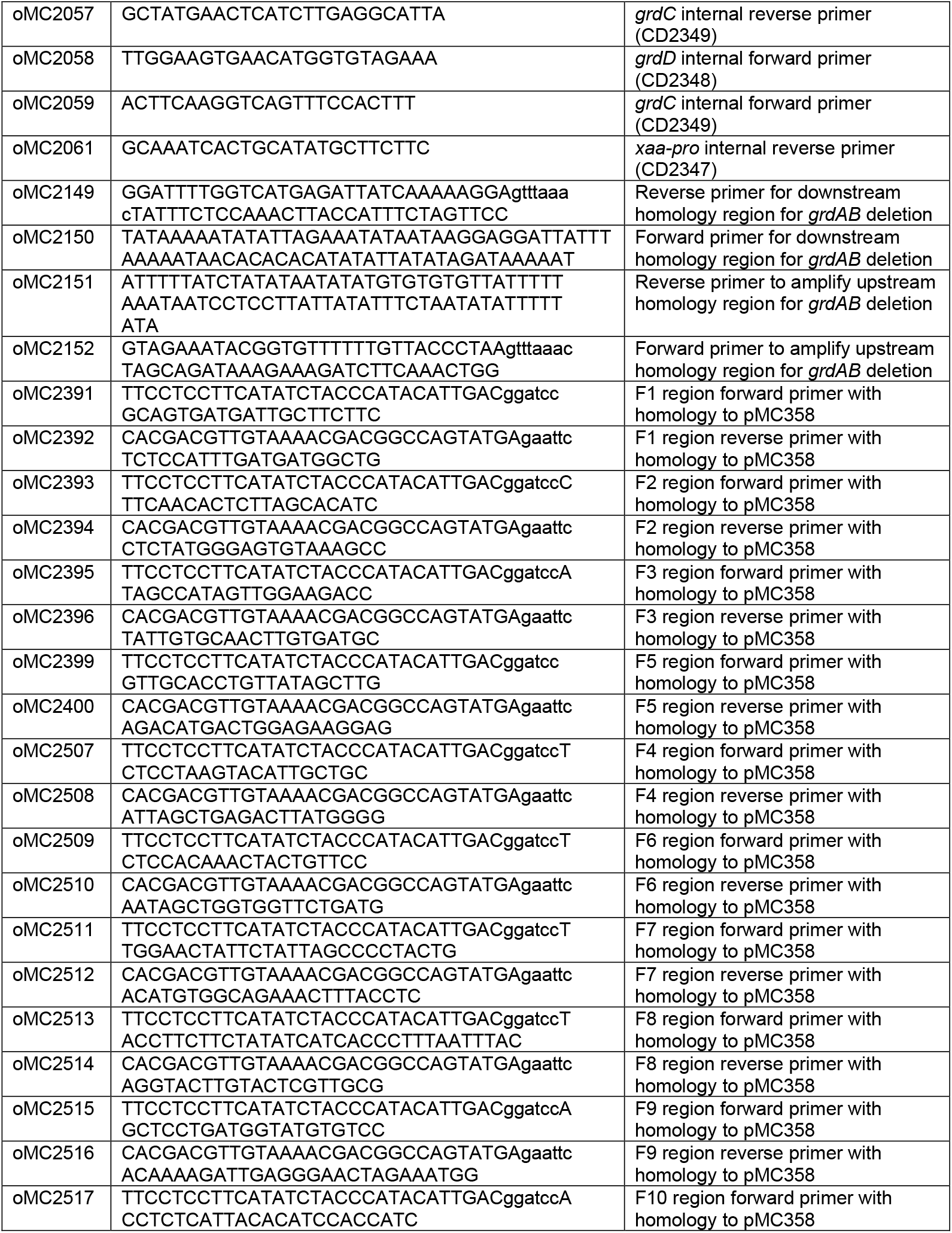
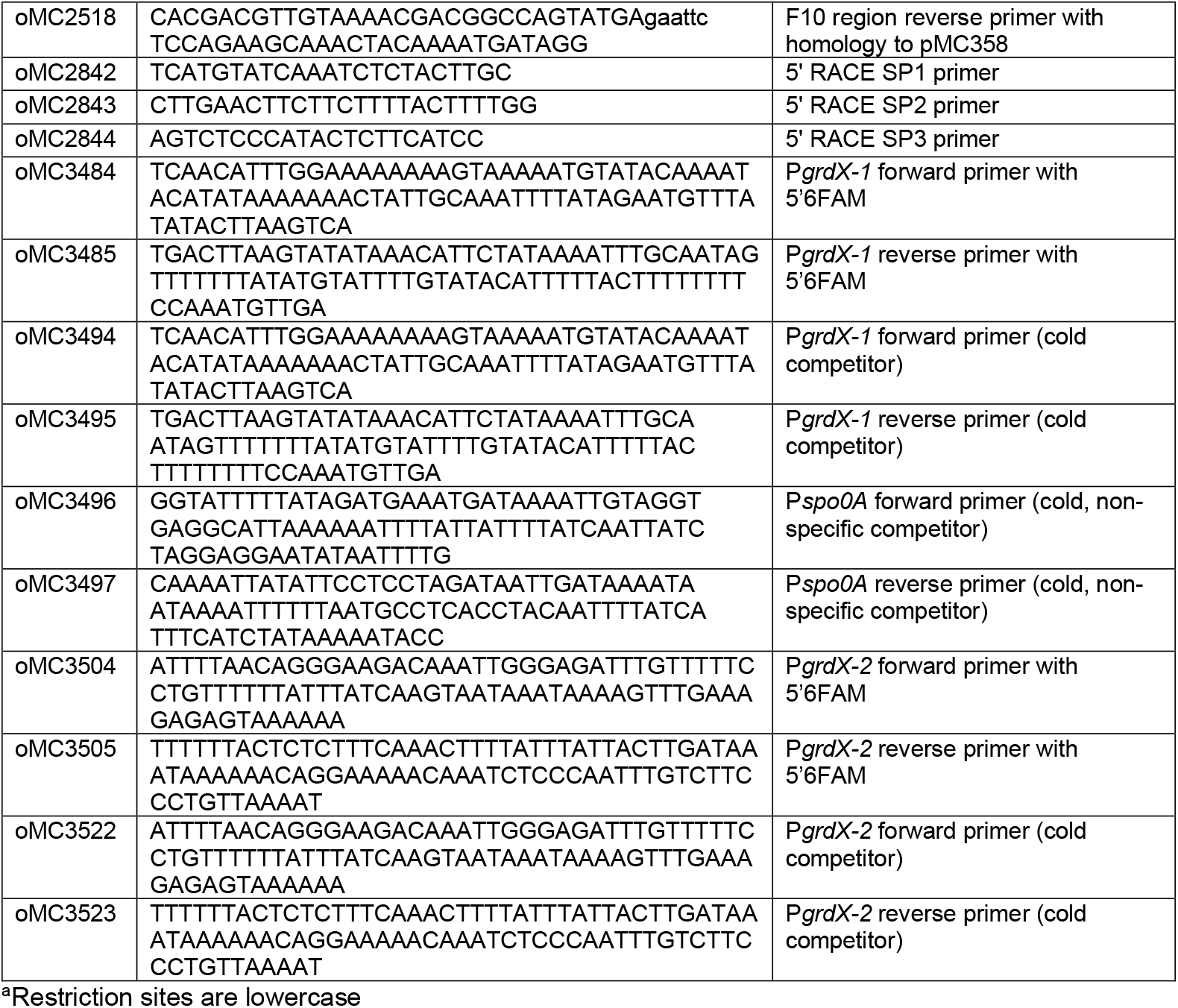
Oligonucleotides.

### Whole Genome Sequencing

Genomic DNA was extracted by Bust n’ Grab protocol (Harju *et al*., 2004). Library prep and paired-end Illumina sequencing (2×151bp) using the Illumina NextSeq2000 platform was performed by SeqCenter/MiGS for WGS, with variant calling service using breseq v0.35.7 (Deatherage and Barrick, 2014)). Geneious Prime 2020.2 was used to perform additional variant calling. A *de novo* assembly was performed using SPAdes to verify the deleted *grdAB* genomic region, and the genome was annotated to the reference sequence (NC_009089.1) using Geneious Prime. (Bankevich *et al*., 2012). Whole genome sequence data for MC1576 (*grdAB*) was deposited in the NCBI SRA database (SAMN23038927).

### Bacterial strains and growth conditions

*Escherichia coli* was grown in lysogeny broth (LB) medium (Teknova) at 37°C and supplemented with 20 µg chloramphenicol ml^-1^ (Sigma-Aldrich) and/or 100 µg ampicillin ml^-1^ (Cayman Chemical Company), when needed. *C. difficile* strains were grown in an anaerobic chamber at 37°C containing 85% N_2_, 10% H_2_, and 5% CO_2_ (Coy Laboratory Products) in brain heart infusion medium supplemented with 2% yeast extract (BHIS; Becton, Dickinson, and Company), unless otherwise noted (Bouillaut *et al*., 2011). *C. difficile* cultures were supplemented with 2-5 µg thiamphenicol ml^-1^ (Sigma-Aldrich) for plasmid selection, as necessary. Counter-selection against *E. coli* post-conjugation was performed by supplementing the medium with 100 µg ml^-1^ kanamycin as previously described (Purcell *et al*., 2012).

Growth in *C. difficile* minimal medium (CDMM) was performed as previously described, but with the addition of 1 µM sodium selenite, 50 µM zinc chloride, and (NH_4_)_2_SO4 at 60.5 µM (Woods *et al*., 2018). As needed, the base CDMM was supplemented with 30 mM glycine, 0.5 µg LL-37 ml^-1^ (Anaspec), or both.

### Sporulation efficiency assays

Sporulation assays were performed as previously described (Childress *et al*., 2016; Edwards and McBride, 2017). Briefly, *C. difficile* strains were cultured in BHIS medium supplemented with 0.1% taurocholate and 0.2% fructose to prevent sporulation, then sub-cultured in BHIS to OD_600_ of 0.5, and 0.25 ml culture plated onto fresh 70:30 sporulation agar (Putnam *et al*., 2013). After 24 h, cells were scraped from the plates and suspended in BHIS medium to an OD_600_ of ∼1.0. The total number of vegetative cells were determined by immediately plating onto BHIS plates. To enumerate spores, cells were treated with ethanol for 15 minutes, then diluted in 1X PBS + 0.1% taurocholate, and plated onto BHIS supplemented with 0.1% taurocholate. CFU were enumerated after at least 36 h of growth, and sporulation frequencies were calculated as the ratio of ethanol-resistant spores divided by the total cells. A *spo0A* mutant (MC310) was used as a negative control to ensure vegetative cell killing.

### Phase-contrast microscopy

Phase-contrast microscopy, samples were resuspended in BHIS broth and applied to slides on agarose pads as previously described (Edwards, Nawrocki et al. 2014). Phase-contrast microscopy was performed using a Nikon Eclipse Ci-L microscope with a X100 Ph3 oil-immersion objective and images were acquired with a DS-Fi2 camera. Two or more fields of view were captured for each strain and growth condition.

### Quantitative reverse transcription PCR (qRT-PCR)

*C. difficile* strains were grown in CDMM as described above. Cells were harvested from CDMM cultures at mid-log phase (defined as an OD_600_ of ∼0.2), pelleted, and ice cold ethanol:acetone (1:1) added before storing at -80°C (Dineen *et al*., 2010). RNA was extracted and treated with DNase I (Ambion) before synthesis of cDNA using random hexamers (Bioline; Edwards, Nawrocki et al. 2014). Quantitative reverse-transcription PCR (qRT-PCR) was performed in technical triplicate with 50 ng cDNA, using the SensiFAST SYBR & Fluorescein Kit (Bioline), on a Roche Lightcycler 96. Reactions without reverse transcriptase were included for all samples to check for contaminating genomic DNA. The comparative cycle threshold method (Schmittgen and Livak, 2008) was used to calculate the results by normalizing expression to the internal control transcript, *rpoC*. A two-tailed Student’s *t* test was used to compare the transcripts in the *grdAB* mutant to the 630Δ*erm* parent strain, as indicated in the figure legend.

### Spore germination assays

*C. difficile* spores were prepared and purified from sporulation agar as previously described (Nawrocki *et al*., 2018). Briefly, spores were scraped and resuspended from plates in 1x PBS + 1% BSA, frozen at -80°C, and repeatedly purified by centrifugation over a 12 ml 50% sucrose bed volume until preparations reached a purity of greater than 95%, as assessed by phase-contrast microscopy. Spores were then washed and resuspended in 1x PBS + 1% BSA and used for germination assays at a starting density of ∼0.3 (OD_600_). Germination frequency was measured by the loss of optical density (OD_600_) after the addition of 5 mM germinant (taurocholate, Sigma-Aldrich), as previously described (Sorg and Sonenshein, 2008; Sorg and Sonenshein, 2009).

### Animal Studies

The Syrian Golden hamster model was used for infection studies with the *grdAB* mutant and parent strain 630Δ*erm* as previously described (Nawrocki *et al*., 2018). Hamsters were housed individually in a sterile ABSL-2 facility within the Emory University Division of Animal Resources. Animals were provided irradiated rodent chow and sterile water, ad libitum. Briefly, 6 -8 week old male and female hamsters from Charles River Laboratories were orally gavaged with 30 mg/kg clindamycin to induce susceptibility to *C. difficile* seven days prior to inoculation by gavage with ∼5000 *C. difficile* spores. Independent *C. difficile* spore inocula were prepared and enumerated prior to each experiment, as previously described (Edwards and McBride, 2016). Prior to infection, spores were heated for 20 min at 55°C and cooled to room temperature before inoculation. Two independent experiments were performed with cohorts of 5-6 hamsters per *C. difficile* strain. Negative control animals were treated with clindamycin and remained free of *C. difficile* throughout the experiments.

Infected animals were monitored for signs of disease, such as wet tail, weight loss, and lethargy and were considered moribund if they lost greater than 15% of their body weight and/or exhibited signs of severe disease or distress. Hamsters that met the moribund criteria were euthanized in accordance with the American Veterinary Medical Association guidelines. Animal survival data (time to morbidity) were analyzed by the Log-rank test. Infected animals were weighed daily and fecal samples were collected for enumeration of CFU. Fecal and cecal contents were plated onto selective TCCFA plates (George *et al*., 1979; Wilson *et al*., 1982) and incubated anaerobically for 48 h prior to enumeration.

### Toxin Quantification

Quantification of TcdA and TcdB toxin was performed using culture supernatant from *C. difficile* strains grown for 24 h in TY medium, pH 7.4 (Garnier and Cole, 1986), as previously described (Edwards *et al*., 2020). Briefly, cultures were pelleted and supernatants were assayed by ELISA in technical duplicate using the tgcBIOMICS toxin A-toxin B kit, per manufacturer’s instructions. A minimum of three biological replicates were performed. *In vivo* cecal toxin was measured similarly from pelleted cecal contents retrieved from hamsters post-mortem.

### Alkaline phosphatase (AP) activity assays

*C. difficile* strains were grown in CDMM broth with 2 µg ml^-1^ thiamphenicol for plasmid maintenance and supplemented with 30 mM glycine, 0.75 µg LL-37 ml^-1^, or both. Samples (1 ml technical duplicates) of *C. difficile* harboring transcriptional fusions of potential *grd* promoter fragments to a *phoZ* reporter gene were harvested at stationary phase, pelleted, and stored at - 20°C until processing. AP assays were performed as previously detailed (Edwards *et al*., 2015; Edwards *et al*., 2019), in the absence of chloroform. Assay results from technical duplicates were averaged, and the results provided as the mean and standard deviation for at least three biological replicates.

### 5’ RACE

RNA was extracted from cells collected as described above for qRT-PCR. After treating the RNA samples with DNase I, the samples were used as template to generate cDNA using the Roche 5’/3’ RACE (rapid amplification of cDNA ends) second-generation kit, following the manufacturer’s protocol. Primers used are listed in **Table 2**. The resulting PCR products were purified and cloned into pCR2.1 (Invitrogen TOPO TA cloning kit) for sequencing (Genewiz).

### Electrophoretic mobility shift assays (EMSAs)

Recombinant *C. difficile* His_6_-ClnR was produced by GenScript (Piscataway, NJ). Quantitative EMSAs were performed as previously described (Woods, Edwards et al. 2018), with the following noted exceptions. Briefly, 5’-fluorescein (5’-6FAM)-labeled DNA probes and unlabeled specific and nonspecific competitors were produced by annealing complementary oligos (**Table 2**) synthesized by IDT (Coralville, IA) in Duplex Buffer (100 mM potassium acetate, 30 mM HEPES, pH 7.5). The resulting DNA probes were diluted in nuclease-free dH_2_O to the working concentration. The 5’-6FAM-labeled DNA probes (10 ng per reaction) were incubated for 30 min at 37°C with the indicated concentration of His_6_-ClnR (0 – 32 µM) and 10 mM Tris, pH 7.4, 10 mM MgCl_2_, 100 mM KCl, 7.5% glycerol, 50 ng salmon sperm DNA, and 2 mM DTT. Competitions assays were performed with 100 ng (10X) and 1 µg (100X) excess unlabeled specific (P*grdX*) or nonspecific (P*spo0A*) DNA. Reactions were subsequently separated on a pre-run 5% TBE gel (Bio-Rad; Hercules, CA). Afterwards, the gel was rinsed with dH_2_O and imaged on a Typhoon Phosphorimager (GE Lifesciences). Densitometry of free and bound DNA species were quantitated with Image Lab (Bio-Rad; Hercules, CA). The apparent K*d* value of P*grdX*-1 was calculated by non-linear regression using a cooperative binding equation with GraphPad Prism 9.4.0 as previously described (Mercante *et al*., 2006; Woods *et al*., 2018).

## REFERENCES

Abt, M.C., McKenney, P.T., and Pamer, E.G. (2016) Clostridium difficile colitis: pathogenesis and host defence. Nat Rev Microbiol 14: 609–620.

Andreesen, J.R. (2004) Glycine reductase mechanism. Current Opinion in Chemical Biology 8: 454–461.

Antunes, A., Camiade, E., Monot, M., Courtois, E., Barbut, F., Sernova, N.V., et al. (2012) Global transcriptional control by glucose and carbon regulator CcpA in Clostridium difficile. Nucleic acids research 40: 10701–18.

Bankevich, A., Nurk, S., Antipov, D., Gurevich, A.A., Dvorkin, M., Kulikov, A.S., et al. (2012) SPAdes: A New Genome Assembly Algorithm and Its Applications to Single-Cell Sequencing. Journal of Computational Biology 19: 455–477.

Battaglioli, E.J., Hale, V.L., Chen, J., Jeraldo, P., Ruiz-Mojica, C., Schmidt, B.A., et al. (2018) Clostridioides difficile uses amino acids associated with gut microbial dysbiosis in a subset of patients with diarrhea. Sci Transl Med 10.

Berges, M., Michel, A.-M., Lassek, C., Nuss, A.M., Beckstette, M., Dersch, P., et al. (2018) Iron Regulation in Clostridioides difficile. Front Microbiol 9: 3183.

Bouillaut, L., Dubois, T., Francis, M.B., Daou, N., Monot, M., Sorg, J.A., et al. (2019) Role of the global regulator Rex in control of NAD+ -regeneration in Clostridioides (Clostridium) difficile. Mol Microbiol 111: 1671–1688.

Bouillaut, L., Dubois, T., Sonenshein, A.L., and Dupuy, B. (2015) Integration of metabolism and virulence in Clostridium difficile. Research in microbiology 166: 375–83.

Bouillaut, L., McBride, S.M., and Sorg, J.A. (2011) Genetic manipulation of Clostridium difficile. Current protocols in microbiology Chapter 9: Unit 9A 2.

Bouillaut, L., Self, W.T., and Sonenshein, A.L. (2013) Proline-dependent regulation of Clostridium difficile Stickland metabolism. Journal of bacteriology 195: 844–54.

Childress, K.O., Edwards, A.N., Nawrocki, K.L., Woods, E.C., Anderson, S.E., and McBride, S.M. (2016) The Phosphotransfer Protein CD1492 Represses Sporulation Initiation in Clostridium difficile. Infection and immunity.

Collery, M.M., Kuehne, S.A., McBride, S.M., Kelly, M.L., Monot, M., Cockayne, A., et al. (2017) What’s a SNP between friends: The influence of single nucleotide polymorphisms on virulence and phenotypes of Clostridium difficile strain 630 and derivatives. Virulence 8: 767–781.

Deatherage, D.E., and Barrick, J.E. (2014) Identification of Mutations in Laboratory-Evolved Microbes from Next-Generation Sequencing Data Using breseq. In Engineering and Analyzing Multicellular Systems. Sun, L., and Shou, W. (eds). Springer New York, New York, NY. pp. 165–188 http://link.springer.com/10.1007/978-1-4939-0554-6_12. Accessed October 26, 2022.

Dineen, S.S., McBride, S.M., and Sonenshein, A.L. (2010) Integration of metabolism and virulence by Clostridium difficile CodY. Journal of bacteriology 192: 5350–62.

Dineen, S.S., Villapakkam, A.C., Nordman, J.T., and Sonenshein, A.L. (2007) Repression of Clostridium difficile toxin gene expression by CodY. Molecular microbiology 66: 206–19.

Edwards, A.N., Anjuwon-Foster, B.R., and McBride, S.M. (2019) RstA Is a Major Regulator of Clostridioides difficile Toxin Production and Motility. mBio 10.

Edwards, A.N., Krall, E.G., and McBride, S.M. (2020) Strain-Dependent RstA Regulation of Clostridioides difficile Toxin Production and Sporulation. J Bacteriol 202 https://journals.asm.org/doi/10.1128/JB.00586-19. Accessed September 18, 2021.

Edwards, A.N., and McBride, S.M. (2016) Isolating and Purifying Clostridium difficile Spores. Methods Mol Biol 1476: 117–28.

Edwards, A.N., and McBride, S.M. (2017) Determination of the in vitro Sporulation Frequency of Clostridium difficile. Bio-protocol 7.

Edwards, A.N., Nawrocki, K.L., and McBride, S.M. (2014) Conserved oligopeptide permeases modulate sporulation initiation in Clostridium difficile. Infect Immun 82: 4276–4291.

Edwards, A.N., Pascual, R.A., Childress, K.O., Nawrocki, K.L., Woods, E.C., and McBride, S.M. (2015) An alkaline phosphatase reporter for use in Clostridium difficile. Anaerobe 32: 98–104.

Fletcher, J.R., Pike, C.M., Parsons, R.J., Rivera, A.J., Foley, M.H., McLaren, M.R., et al. (2021) Clostridioides difficile exploits toxin-mediated inflammation to alter the host nutritional landscape and exclude competitors from the gut microbiota. Nat Commun 12: 462.

Garnier, T., and Cole, S.T. (1986) Characterization of a bacteriocinogenic plasmid from Clostridium perfringens and molecular genetic analysis of the bacteriocin-encoding gene. J Bacteriol 168: 1189–1196.

George, W.L., Rolfe, R.D., Sutter, V.L., and Finegold, S.M. (1979) Diarrhea and colitis associated with antimicrobial therapy in man and animals. The American journal of clinical nutrition 32: 251–7.

Harju, S., Fedosyuk, H., and Peterson, K.R. (2004) Rapid isolation of yeast genomic DNA: Bust n’ Grab. BMC biotechnology 4: 8.

Hofmann, J.D., Otto, A., Berges, M., Biedendieck, R., Michel, A.M., Becher, D., et al. (2018) Metabolic Reprogramming of Clostridioides difficile During the Stationary Phase With the Induction of Toxin Production. Frontiers in microbiology 9: 1970.

Hussain, H.A., Roberts, A.P., and Mullany, P. (2005) Generation of an erythromycin-sensitive derivative of Clostridium difficile strain 630 (630Δerm) and demonstration that the conjugative transposon Tn916ΔE enters the genome of this strain at multiple sites. Journal of medical microbiology 54: 137–141.

Jackson, S., Calos, M., Myers, A., and Self, W.T. (2006) Analysis of proline reduction in the nosocomial pathogen Clostridium difficile. Journal of bacteriology 188: 8487–95.

Johnstone, M.A., and Self, W.T. (2022) D -Proline Reductase Underlies Proline-Dependent Growth of Clostridioides difficile. J Bacteriol e00229–22.

Karasawa, T., Ikoma, S., Yamakawa, K., and Nakamura, S. (1995) A defined growth medium for Clostridium difficile. Microbiology 141: 371–375.

Karlsson, S., Burman, L.G., and Akerlund, T. (2008) Induction of toxins in Clostridium difficile is associated with dramatic changes of its metabolism. Microbiology 154: 3430–6.

Lee, C.D., Rizvi, A., Edwards, A.N., DiCandia, M.A., Vargas Cuebas, G.G., Monteiro, M.P., and McBride, S.M. (2022) Genetic mechanisms governing sporulation initiation in Clostridioides difficile. Current Opinion in Microbiology 66: 32–38.

McBride, S.M., and Sonenshein, A.L. (2011) Identification of a genetic locus responsible for antimicrobial peptide resistance in Clostridium difficile. Infect Immun 79: 167–176.

Mercante, J., Suzuki, K., Cheng, X., Babitzke, P., and Romeo, T. (2006) Comprehensive Alanine-scanning Mutagenesis of Escherichia coli CsrA Defines Two Subdomains of Critical Functional Importance. Journal of Biological Chemistry 281: 31832–31842.

Nawrocki, K.L., Edwards, A.N., Daou, N., Bouillaut, L., and McBride, S.M. (2016) CodY-Dependent Regulation of Sporulation in Clostridium difficile. Journal of bacteriology 198: 2113–30.

Nawrocki, K.L., Wetzel, D., Jones, J.B., Woods, E.C., and McBride, S.M. (2018) Ethanolamine is a valuable nutrient source that impacts Clostridium difficile pathogenesis. Environmental microbiology 20: 1419–1435.

Neumann-Schaal, M., Hofmann, J.D., Will, S.E., and Schomburg, D. (2015) Time-resolved amino acid uptake of Clostridium difficile 630Deltaerm and concomitant fermentation product and toxin formation. BMC microbiology 15: 281.

Neumann-Schaal, M., Jahn, D., and Schmidt-Hohagen, K. (2019) Metabolism the Difficile Way: The Key to the Success of the Pathogen Clostridioides difficile. Frontiers in microbiology 10: 219.

Paredes-Sabja, D., Shen, A., and Sorg, J.A. (2014) Clostridium difficile spore biology: sporulation, germination, and spore structural proteins. Trends in microbiology 22: 406–416.

Peltier J, H.A. (2020) Type I toxin-antitoxin systems contribute to mobile genetic elements maintenance in Clostridioides difficile and can be used as a counter-selectable marker for chromosomal manipulation. BioRxiv.

Purcell, E.B., McKee, R.W., McBride, S.M., Waters, C.M., and Tamayo, R. (2012) Cyclic diguanylate inversely regulates motility and aggregation in Clostridium difficile. Journal of bacteriology 194: 3307–16.

Putnam, E.E., Nock, A.M., Lawley, T.D., and Shen, A. (2013) SpoIVA and SipL are Clostridium difficile spore morphogenetic proteins. Journal of bacteriology 195: 1214–25.

Sandhu, B.K., and McBride, S.M. (2018) .Clostridioides difficile. Trends in microbiology 26: 1049–1050.

Saujet, L., Monot, M., Dupuy, B., Soutourina, O., and Martin-Verstraete, I. (2011) The key sigma factor of transition phase, SigH, controls sporulation, metabolism, and virulence factor expression in Clostridium difficile. Journal of bacteriology 193: 3186–96.

Schmittgen, T.D., and Livak, K.J. (2008) Analyzing real-time PCR data by the comparative C(T) method. Nature protocols 3: 1101–8.

Sorg, J.A., and Sonenshein, A.L. (2008) Bile salts and glycine as cogerminants for Clostridium difficile spores. Journal of bacteriology 190: 2505–12.

Sorg, J.A., and Sonenshein, A.L. (2009) Chenodeoxycholate is an inhibitor of Clostridium difficile spore germination. Journal of bacteriology 191: 1115–7.

Soutourina, O., Dubois, T., Monot, M., Shelyakin, P.V., Saujet, L., Boudry, P., et al. (2020) Genome-Wide Transcription Start Site Mapping and Promoter Assignments to a Sigma Factor in the Human Enteropathogen Clostridioides difficile. Front Microbiol 11: 1939.

Soutourina, O.A., Monot, M., Boudry, P., Saujet, L., Pichon, C., Sismeiro, O., et al. (2013) Genome-wide identification of regulatory RNAs in the human pathogen Clostridium difficile. PLoS genetics 9: e1003493.

Stickland, L.H. (1934) Studies in the metabolism of the strict anaerobes (genus Clostridium): The chemical reactions by which Cl. sporogenes obtains its energy. The Biochemical journal 28: 1746–59.

Stickland, L.H. (1935) Studies in the metabolism of the strict anaerobes (genus Clostridium): The oxidation of alanine by Cl. sporogenes. IV. The reduction of glycine by Cl. sporogenes. The Biochemical journal 29: 889–98.

Thomas, C.M., and Smith, C.A. (1987) Incompatibility group P plasmids: genetics, evolution, and use in genetic manipulation. Annual Reviews in Microbiology 41: 77–101.

Wilson, K.H., Kennedy, M.J., and Fekety, F.R. (1982) Use of sodium taurocholate to enhance spore recovery on a medium selective for Clostridium difficile. Journal of clinical microbiology 15: 443–6.

Woods, E.C., Edwards, A.N., Childress, K.O., Jones, J.B., and McBride, S.M. (2018) The C. difficile clnRAB operon initiates adaptations to the host environment in response to LL-37. PLoS pathogens 14: e1007153.

Yanisch-Perron, C., Vieira, J., and Messing, J. (1985) Improved M13 phage cloning vectors and host strains: nucleotide sequences of the M13mp18 and pUC19 vectors. Gene 33: 103–19.

