## Supplemental for "Glycine fermentation by *C. difficile* promotes virulence, spore formation, and is induced by host cathelicidin"

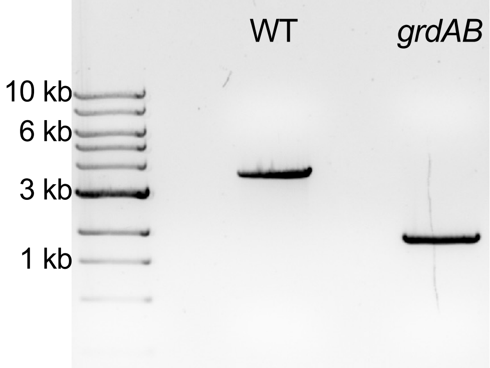


**Figure S1. Confirmation of the *grdAB* mutant by PCR.** Amplification of *grdA*-*grdB* region from genomic DNA of strains 630Δ*erm* and the *grdAB* deletion mutant (MC1576) using flanking primers (oMC2052/oMC2057). Expected sizes are 3.2 kb for the parent strain (WT, 630Δ*erm*) and 1.8 kb for the *grdAB* mutant. NEB 1 kb ladder shown.

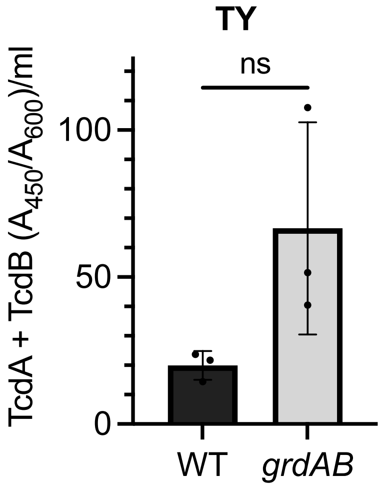


**Figure S2. The *grdAB* mutant displays variable toxin production.** *C. difficile* 630Δ*erm* (WT) and *grdAB* (MC1576) mutant total toxin (TcdA + TcdB) was quantified by ELISA from supernatants after growth for 24 h in TY medium. The means and SD for a minimum of three biological replicates are shown. No statistical significance was found by Student's *t* test.

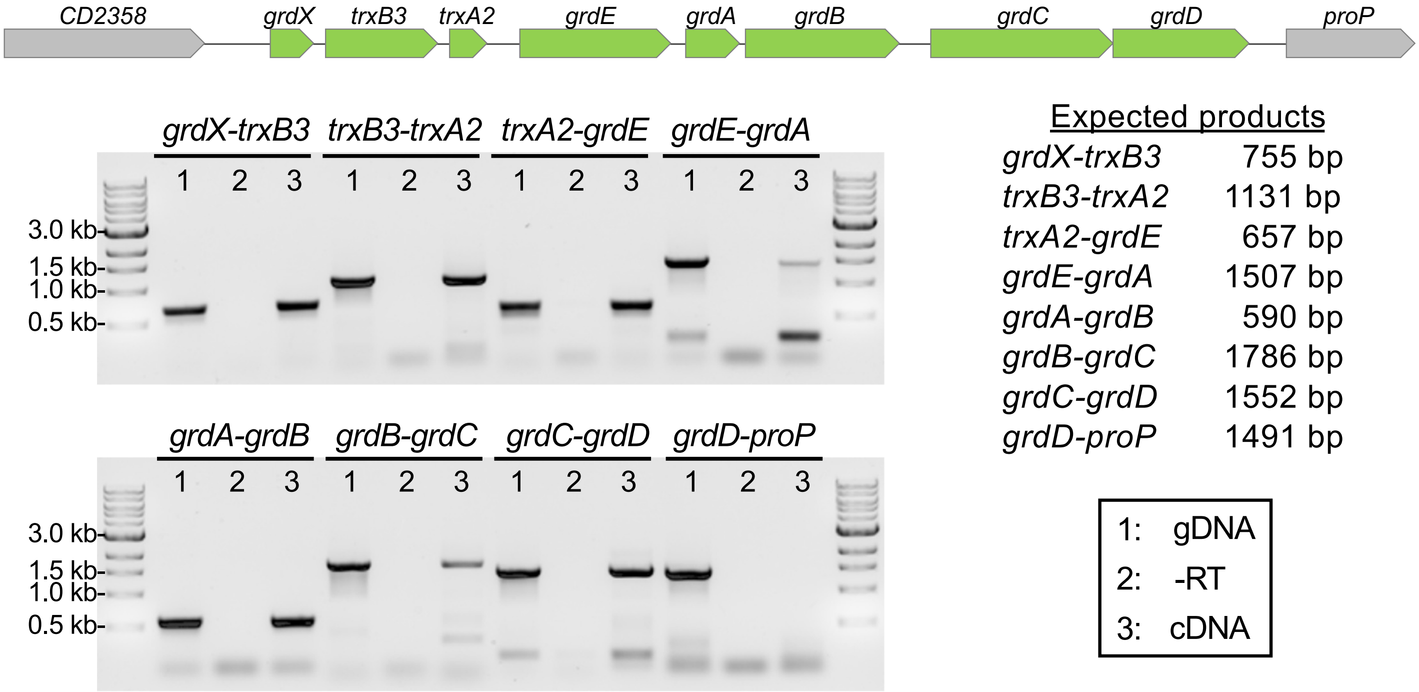


**Figure S3. Analysis of transcriptional units within the *grd* gene cluster*.*** Transcriptional units were examined by PCR using designed to generate products if adjacent *grd* open reading frames are produced on the same transcript. Templates for each primer pair were as follows: 630Δ*erm* genomic DNA (gDNA, positive control), cDNA generated from *C. difficile* grown on 70:30 sporulation medium for six hours, and mock cDNA reaction without reverse transcriptase (negative control). Genes encoded in the *grd* operon are colored green. Primer pairs include: *grdX-trxB3,* oMC2046/2049; *trxB3-trxA2,* oMC2048/2051; *trxA2-grdE,* oMC2050/2053; *grdE-grdA,* oMC2052/oMC1384; *grdA-grdB,* oMC1383/oMC2055; *grdB-grdC,* oMC2054/2057; *grdC-grdD,* oMC2056/2059; *grdD-CD2347,* oMC2058/2061.

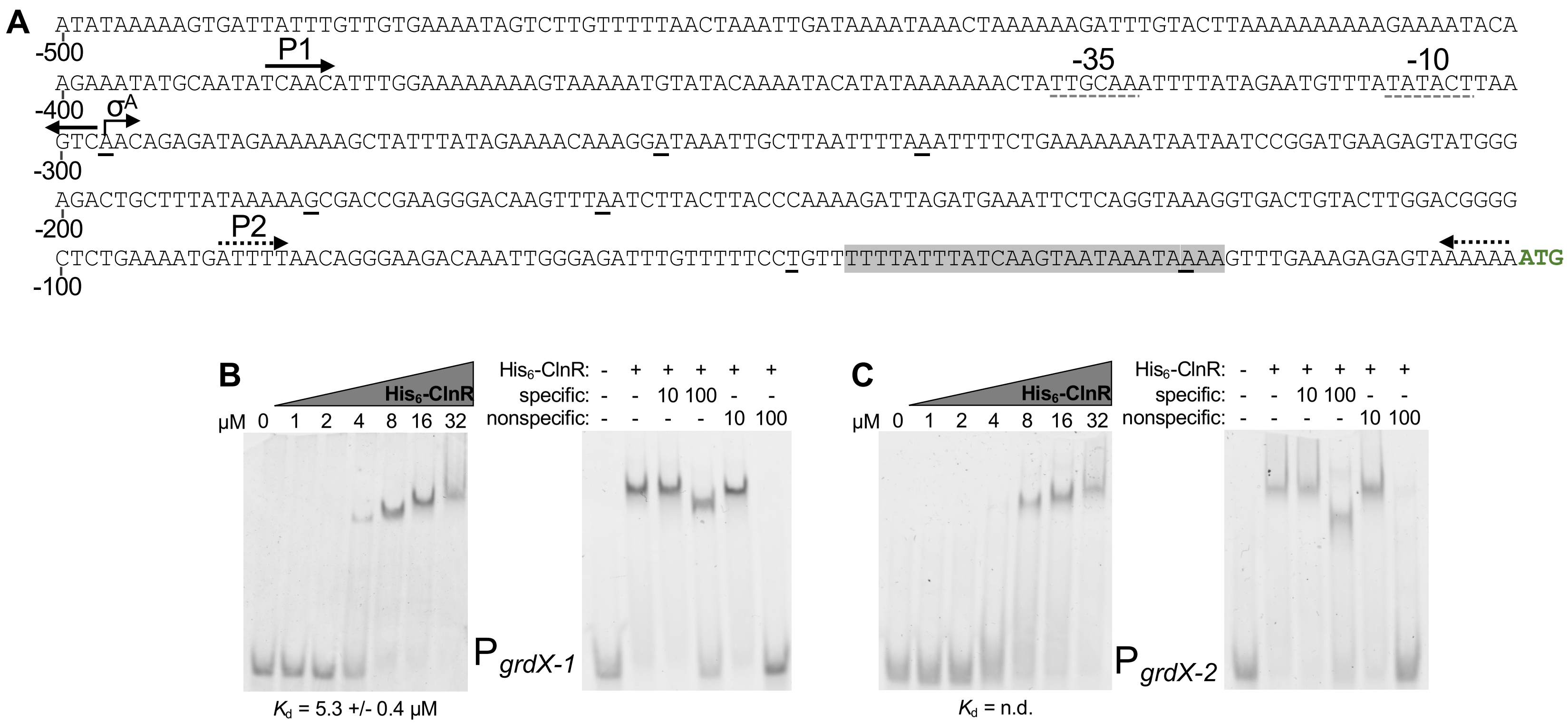


**Figure S4. Transcriptional start site mapping and ClnR binding to P*grdX*.** A) Schematic of the *grdX* promoter region. Transcriptional start sites identified by 5' RACE are underlined in black, a predicted SigA promoter is marked with gray dashed lines, and an inverted repeat is shaded. B) and C) Electrophoretic mobility shift assays were performed using N-terminally His-tagged ClnR and fluorescein-labeled DNA. Binding of ClnR to the B) P*grdX-1* or C) P*grdX-2* DNA probe with increasing concentrations of ClnR protein. Competitive EMSAs performed with the addition of unlabeled specific or unlabeled nonspecific (P*spo0A*) DNA at either 10x or 100x the concentration of labeled probe. n.d.: not determined due to instability

**Figure S5. DNA cloning and vector details**

pMC950: A 1.082 kb PCR product containing a 569 bp and 545 bp region of upstream and downstream flanking homology to *grdAB* was generated by splicing by overlap PCR with fragments amplified from primers oMC2149/2150 and oMC2151/2152, respectively. This PCR product was Gibson assembled into pMSR at the PmeI site.

pMC951: A 882 bp PCR fragment of the comprising the *CD2358-grdX* region (F1) was amplified with primers oMC2391/2392 and Gibson assembled into pMC358 at the BamHI and EcoRI sites.

pMC952: A 885 bp PCR fragment of the comprising the *grdX-trxB3* region (F2) was amplified with primers oMC2393/2394 and Gibson assembled into pMC358 at the BamHI and EcoRI sites.

pMC953: A 879 bp PCR fragment of the comprising the *trxB3-trxA2* region (F3) was amplified with primers oMC2395/2396 and Gibson assembled into pMC358 at the BamHI and EcoRI sites.

pMC954: A 961 bb PCR fragment of the comprising the *trxA2-grdE* 5’ region (F4) was amplified with primers oMC2507/2508 and Gibson assembled into pMC358 at the BamHI and EcoRI sites.

pMC955: A 916 bp PCR fragment of the comprising the *grdE* 5’-*grdE* 3’ region (F5) was amplified with primers oMC2399/2400 and Gibson assembled into pMC358 at the BamHI and EcoRI sites.

pMC956: A 910 bp PCR fragment of the comprising the *grdE* 3’-*grdB* 5’ region (F6) was amplified with primers oMC2509/2510 and Gibson assembled into pMC358 at the BamHI and EcoRI sites.

pMC957: A 1094 bp PCR fragment of the comprising the *grdB* 5’-*grdB* 3’ region (F7) was amplified with primers oMC2511/2512 and Gibson assembled into pMC358 at the BamHI and EcoRI sites.

pMC958: A 965 bp PCR fragment of the comprising the *grdB* 3’-*grdC* 5’ region (F8) was amplified with primers oMC2513/2514 and Gibson assembled into pMC358 at the BamHI and EcoRI sites.

pMC959: A 1057 bp PCR fragment of the comprising the *grdC* 5’-*grdC* 3’ region (F9) was amplified with primers oMC2515/2516 and Gibson assembled into pMC358 at the BamHI and EcoRI sites.

pMC960: A 983 bp PCR fragment of the comprising the *grdC* 3’-*grdD* region (F10) was amplified with primers oMC2517/2518 and Gibson assembled into pMC358 at the BamHI and EcoRI sites.
